# Graded changes in local functional connectivity of the cerebral cortex in young people with depression

**DOI:** 10.1101/2023.10.23.563507

**Authors:** Alec J. Jamieson, Christopher G. Davey, Jesus Pujol, Laura Blanco-Hinojo, Ben J. Harrison

**Author notes:** **Corresponding author:** Dr. Alec Jamieson. Melbourne Neuropsychiatry Centre, Department of Psychiatry, The University of Melbourne. Level 3, 161 Barry St. Carlton, VIC 3053. |.

## Abstract

Major depressive disorder (MDD) is marked by significant changes to the coupling of spontaneous neural activity within various brain regions. However, many methods for assessing this local connectivity use fixed or arbitrary neighborhood sizes, resulting in a decreased capacity to capture smooth changes to the spatial gradient of local correlations. A newly developed method sensitive to classical anatomo-functional boundaries, Iso-Distant Average Correlation (IDAC), was therefore used to examine depression associated alterations to the local functional connectivity of the brain. One-hundred and forty-five adolescents and young adults with MDD and 95 healthy controls underwent a resting-state fMRI scan. Whole-brain functional connectivity maps of intracortical neural activity within iso-distant local areas (5-10mm, 15-20mm, and 25-30mm) were generated to characterize local fMRI signal similarities. Across all spatial distances, MDD participants demonstrated greater local functional connectivity of the bilateral posterior hippocampus, retrosplenial cortex, dorsal insula, fusiform gyrus, and supplementary motor area. Additionally, in the short and medium range connections there were depression associated alterations in the midcingulate (15-20mm and 25-30mm) and subgenual anterior cingulate (15-20mm). Our study identified increased synchrony of the neural activity in several regions commonly implicated in the neurobiology of depression; however, a subset of identified effects was dependent on the spatial distance under consideration. Longitudinal examination of these effects will clarify whether these differences are also found in other age groups and if this synchrony is additionally altered by continued disease progression.

## Introduction

Major depressive disorder (MDD) is a highly prevalent and disabling mental health condition characterized by pervasive symptoms of sadness, hopelessness, and anhedonia (American Psychiatric Association and Association, 2013). Depression often first arises during adolescence (Gore *et al*., 2011, Thapar *et al*., 2012), with early onset being a strong predictor of recurrent depression (Fergusson *et al*., 2005, Klein *et al*., 2009) as well as greater reductions in quality of life and wellbeing into adulthood (Zisook *et al*., 2007). Early adolescence is marked by increased use of maladaptive emotional regulation strategies which in turn are associated with exacerbated mental health issues (Cracco *et al*., 2017, Silk *et al*., 2003). One such strategy, known expressive suppression, involves inhibiting the outward expression of emotional behavior (Gross, 2015), has been specifically associated with depressive symptoms severity (Larsen *et al*., 2013). However, there remains limited work examining neurobiological underpinnings of this early stage of the disorder.

Previous neuroimaging research has attempted to uncover the neurobiological underpinnings of MDD by focusing on long-range functional connectivity, examining the temporal correlations of large-scale, spatially distinct brain networks. Within this framework, MDD has been associated with alterations across multiple intrinsic brain networks (Brakowski *et al*., 2017, Li *et al*., 2018), including the default mode (DMN), salience, and central executive networks (Dunlop *et al*., 2019). However, there has been increasing interest in assessing local functional connectivity profiles (Jiang and Zuo, 2016, Zuo *et al*., 2013), as the functional integration of local neuronal groups appears to represent a key organizing principle in the brains of higher vertebrates (Tononi *et al*., 1994). Local functional connectivity measures assess the temporal coherence of the functional magnetic resonance imaging (fMRI) signal between a given voxel and neighboring voxels (Jiang and Zuo, 2016). Measures of this ‘localized synchrony’ provide a framework for examining functional disruptions at rest without a priori constraints, thereby allowing for data-driven identification of regional abnormalities across a range of mental health disorders (Guo *et al*., 2011). Moreover, regional variation in homogeneity has been associated with the hierarchical organization of information processing across the brain, and therefore alterations to these measures may serve as key markers of human brain function (Jiang *et al*., 2015, Jiang and Zuo, 2016) and a sensitive tool for identifying brain alterations across mental health disorders (Canario *et al*., 2021, Wei *et al*., 2018).

Meta-analyses examining regional homogeneity in MDD have revealed inconsistent differences in the directionality of effects and regions implicated (Chen *et al*., 2015, Hao *et al*., 2019, Iwabuchi *et al*., 2015). Altered local connectivity has been noted across regions including the parahippocampal gyrus, insula, and medial prefrontal cortex (Chen *et al*., 2015, Iwabuchi *et al*., 2015). Other local functional connectivity measures, including functional connectivity density and dynamic regional phase synchrony, have illustrated reduced local connectivity in the anterior cingulate cortex, precuneus, hippocampus, thalamus, and insula (Ke *et al*., 2016, Tang *et al*., 2022, Zheng *et al*., 2022). Conversely, recent work examining first-episode MDD observed increased regional homogeneity of the hippocampus and insula but reduced local connectivity of the orbitofrontal cortex (Hao *et al*., 2019). It is likely that the specific method used, including how neighborhood boundaries are defined, as well as chronicity of the disorder, contributes to the inconsistencies observed between studies. Despite these inconsistencies, the observed alterations appear to be localized to regions implicated in affective processing and regulation (Rolls and Grabenhorst, 2008, Uddin *et al*., 2017). Of particular note is the implication of insula dysfunction across measures of local and remote functional connectivity (Iwabuchi *et al*., 2014, Jamieson *et al*., 2022, Manoliu *et al*., 2014, Pastrnak *et al*., 2021), due to its hypothesized role in the integration of autonomic, emotional, and interoceptive stimuli (Sliz and Hayley, 2012) and potential specificity in delineating MDD from bipolar disorder (Pastrnak *et al*., 2021).

We aimed to investigate local functional connectivity alterations in a large sample of adolescents and young adults with MDD, using the recently introduced framework - IsoDistant Average Correlation (IDAC) (Macia *et al*., 2018). Measures of local functional connectivity often use arbitrary and binarized neighborhood boundaries, thereby losing the capacity to describe the rich smooth spatial gradient of local fMRI correlations (Sepulcre *et al*., 2010, Tomasi and Volkow, 2010, Zang *et al*., 2004). IDAC assesses the average temporal correlation of one voxel with all neighboring voxels within different spatial lags, thereby overcoming these limitations (Macia *et al*., 2018). Based on previous findings, we hypothesize that MDD participants would illustrate increased local functional connectivity of the insula, anterior cingulate, and hippocampus and reduced local functional connectivity of the orbitofrontal cortex. Given the developmental nature of our cohort, we additionally investigated whether local connectivity was associated with measures of depressive symptom severity, and emotion processing and regulation.

## Methods

One hundred and forty-nine unmedicated, help-seeking MDD participants were recruited as part of the Youth Depression Alleviation-Combined Treatment and -Augmentation trials, for full details see Davey and colleagues (Davey *et al*., 2019) and Berk and colleagues (Berk *et al*., 2020). MDD participants were between 15 to 25 years of age and were recruited through specialist mental health clinics in the northern and western suburbs of Melbourne, Australia. These participants had a current diagnosis of MDD, as assessed by the Structured Clinical Interview for DSM-IV Axis I Disorders (SCID) (First *et al*., 1997). Depressive symptoms were at least of a moderate level of severity, indicated by a Montgomery-Åsberg Depression Rating Scale (MADRS) score greater than or equal to 20. Exclusion criteria included a lifetime or current diagnosis of a psychotic or bipolar disorder, current treatment with antidepressant medication, or MRI contraindications including pregnancy. One hundred age and sex-matched healthy participants were also recruited through online advertisements. They had no past mental health disorder diagnoses as assessed through SCID criteria. Due to excessive head motion (three MDD participants, four controls; see below) and incidental findings (one MDD participant, one healthy control), a total of four MDD participants and five controls were excluded from further analysis. This resulted in 95 healthy controls and 145 MDD participants in our final sample. This study and consent process was approved by the Melbourne Health Human Research and Ethics Committee. An informed consent form was provided to, and signed by, all participants in this study. For participants under the age of 18, both participant and parental consent were required.

### Image Acquisition

A 3T General Electric Signa Excite system with an eight-channel phased-array head coil was used in combination with ASSET parallel imaging. The functional sequence consisted of a single shot gradient-recalled echo-planar imaging sequence in the steady state (repetition time, 2000 ms; echo time, 35 ms; and pulse angle, 90°) in a 23-cm field-of-view, with a 64 x 64-pixel matrix and a slice thickness of 3.5mm (no gap). Thirty-six interleaved slices were acquired parallel to the anterior-posterior commissure line with a 20° anterior tilt to better cover ventral prefrontal brain regions. The total sequence duration was 8 minutes, corresponding to 240 whole-brain echo-planar imaging volumes. Participants were instructed to keep their eyes closed for the duration of the scan. The first 4 volumes from each run were automatically discarded to allow for signal equilibration. A T1-weighted high-resolution anatomical image was acquired for each participant to assist with functional timeseries co-registration (140 contiguous slices; repetition time, 7.9 ms; echo time, 3 ms; flip angle, 13°; in a 25.6 cm field-of-view, with a 256 x 256-pixel matrix and a slice thickness of 1 mm). To assist with noise reduction and head immobility, all participants used earplugs and had their heads supported with foam-padding inserts.

### Image Preprocessing

Imaging data was transferred to a Unix-based platform that ran MATLAB Version 9.3 (The MathWorks Inc., Natick, USA) and Statistical Parametric Mapping (SPM) Version 12 v7487 (Wellcome Trust Centre for Neuroimaging, London, UK). Preprocessing followed previously reported steps for IDAC analysis (Pujol *et al*., 2023). In brief, for each subject functional MRI images were slice-time corrected, realigned, and co-registered to their corresponding anatomical image with an affine transformation. Spatial normalization occurred through a back-transformation process, as such the individual 3D anatomical images were segmented and registered to MNI space so that the resulting deformation fields could be applied to the IDAC maps (see below). Images were then re-sliced to 3x3x3 mm resolution and smoothed by convolving the image with a 4x4x4 mm full width at half maximum Gaussian kernel. Motion fingerprint (Wilke, 2012) was used to quantify participant head motion. Participants were excluded if movement exceeded a mean total displacement of 3 mm (∼1 native voxel).

### IDAC analysis

IDAC maps describe the pattern of correlation decay propagating from each voxel across the brain. This approach differs from other method for assessing local connectivity by using multi-distance local measures, rather than a single local measure. These multi-distance measures can uniquely detail the rich spatial structure of the cerebral cortex functional connections, as local connectivity is distance-specific to a large extent. This graded change in local functional connectivity provides a more comprehensive characterization of the brain’s functional structure and has been shown to differentiate the human cortex into regions consistent with traditional brain atlases as well as being sensitive to different brain functional states (Macia *et al*., 2018). For a detailed overview of IDAC see Macia and colleagues (Macia *et al*., 2018). Whole-cortex IDAC maps were computed using the mean correlation z-score of each voxel with all neighboring voxels placed at increasingly spaced iso-distant intervals. IDAC maps were calculated for three iso-distant intervals: 5-10 mm, 15-20 mm, and 25-30 mm, and conducted separately for the right and left hemisphere in native space. These three intervals were selected based on previously published work (Macia *et al*., 2018). However, thinner intervals and more numerous intervals could additionally be calculated, providing a richer but more noisy characterization of IDAC curves. Notably, beyond a 30-mm radius, negative correlations may give rise to cancellation phenomena. Covariates included in this analysis were the six rigid body realignment parameters, their first-order derivatives, average white matter, cerebrospinal fluid, and global brain signal. Motion scrubbing using the realignment parameters from preprocessing was additionally conducted to discard motion-affected image volumes (Power *et al*., 2014). For volumes with greater than 0.2 mm of inter-frame motion, the corresponding volume and those immediately preceding and succeeding were discarded. A discrete cosine transform filter was applied for frequencies outside the .01–0.1 Hz interval.

The resulting maps were then normalized to the Montreal Neurological Institute (MNI) space using a back-transformation process to enable group inference. RGB color overlays were used for displaying the values from the three distances simultaneously. For this study, red represented results from 5-10 mm IDAC maps, green from 15-20 mm, and blue from 25-30 mm. Overlaps of these distances are illustrated through the respective secondary colors.

### Statistical Analysis

Between groups analysis were conducted using a mixed model ANOVA (group [healthy control and MDD] by distance [5–10, 15–20, and 25–30 mm]). Following examination of the Effect of Group F contrast, post hoc t-tests were performed to examine group differences in both directions for each of the three IDAC maps. This resulted in six primary contrasts of interest. Accordingly, we adjusted the false-discovery error rate (FDR) corrected threshold to .008 (0.05/6). Thus, all results displayed were estimated with a whole-brain, FDR corrected threshold of *p* < .008, *k* > 10 voxels. Secondary level covariates included age, gender, and handedness.

Given the regions implicated in the between groups comparison, we then examined whether local functional connectivity changes were associated with the scores from the Emotional Regulation Questionnaire (ERQ) (Gross and John, 2003). The instrument assesses the two means by which people attempt to regulate their emotions: by expressive suppression (‘Suppression’) and by cognitive reappraisal (‘Reappraisal’). In addition to the aforementioned covariates, MADRS scores were included in these analyses to determine whether these effects were independent of depressive symptom severity. Analyses of between-group differences for clinical and demographic characteristics were calculated in SPSS version 27 (IBMCorp., Armonk, NY). Comparisons were adjusted for multiple comparisons using the Holm-Bonferroni correction (Holm, 1979) to determine significance (*p* < .05).

## Results

### Group Differences Between Demographic and Clinical Characteristics

Demographic and clinical characteristics of the sample are reported in Table 1. Healthy controls and MDD patients differed significantly on MADRS symptoms (*t*(233.5) = - 58.80, *p* < .001), ERQ Suppression (*t*(241) = -5.94, *p* < .001) and Reappraisal scores (*t*(216.6) = 10.60, *p* < .001).

**Table 1.** Comparison of Characteristics Between Healthy Controls and Major Depressive Disorder Patients.

| Characteristics | Healthy Controls<br>(*N* = 95) |  | MDD<br>(*N* = 145) |  | Hedges'<br>g or<br>Cramer's<br>V | *p* |
| --- | --- | --- | --- | --- | --- | --- |
|  | Mean or<br>N | *SD* or<br>Percentage | Mean or<br>N | *SD* or<br>Percentage |  |  |
| Age (years) | 20.1 | 2.9 | 19.9 | 2.8 | .07 | 1.00 |
| Female | 52 | 54.7 | 81 | 54.4 | .01 | 1.00 |
| Handedness (Right) | 78 | 82.1 | 139 | 92.7 | .18 | .095 |
| Ethnicity |  |  |  |  | .23 | 1.00 |
| *Caucasian* | 70 | 73.7 | 122 | 81.9 |  |  |
| *Black or African* | 1 | 1.1 | 1 | 0.1 |  |  |
| *Asian* | 19 | 20.0 | 23 | 15.4 |  |  |
| *South American* | 3 | 3.3 | 1 | 0.1 |  |  |
| *Indian* | 1 | 1.1 | 1 | 0.1 |  |  |
| *Middle Eastern* | 1 | 1.1 | 2 | 0.1 |  |  |
| MADRS score | 2.13 | 2.8 | 33.0 | 5.4 | -6.76 | <.001 |
| ERQ Suppression score | 14.35 | 4.7 | 17.79 | 4.2 | -.78 | <.001 |
| ERQ Reappraisal score | 30.76 | 6.2 | 22.0 | 6.9 | 1.02 | <.001 |
| Age of MDD onset | - | - | 15.5 | 2.7 | - | - |

### Functional Structure of the Cerebral Cortex

To ensure that our results are consistent with previously illustrated anatomo-functional boundaries (Macia *et al*., 2018, Pujol *et al*., 2019, Pujol *et al*., 2023), whole-brain maps were generated across the three distances for the healthy controls (Figure 1). Consistent with previous effects, the visual association cortex demonstrated high connectivity across all local distance ranges. The bilateral inferior parietal lobules, including the angular and supramarginal gyri, demonstrated high connectivity at short and medium distances. Moreover, the prefrontal cortex was comprised of high connectivity in the long and medium distances.

**Figure 1.**
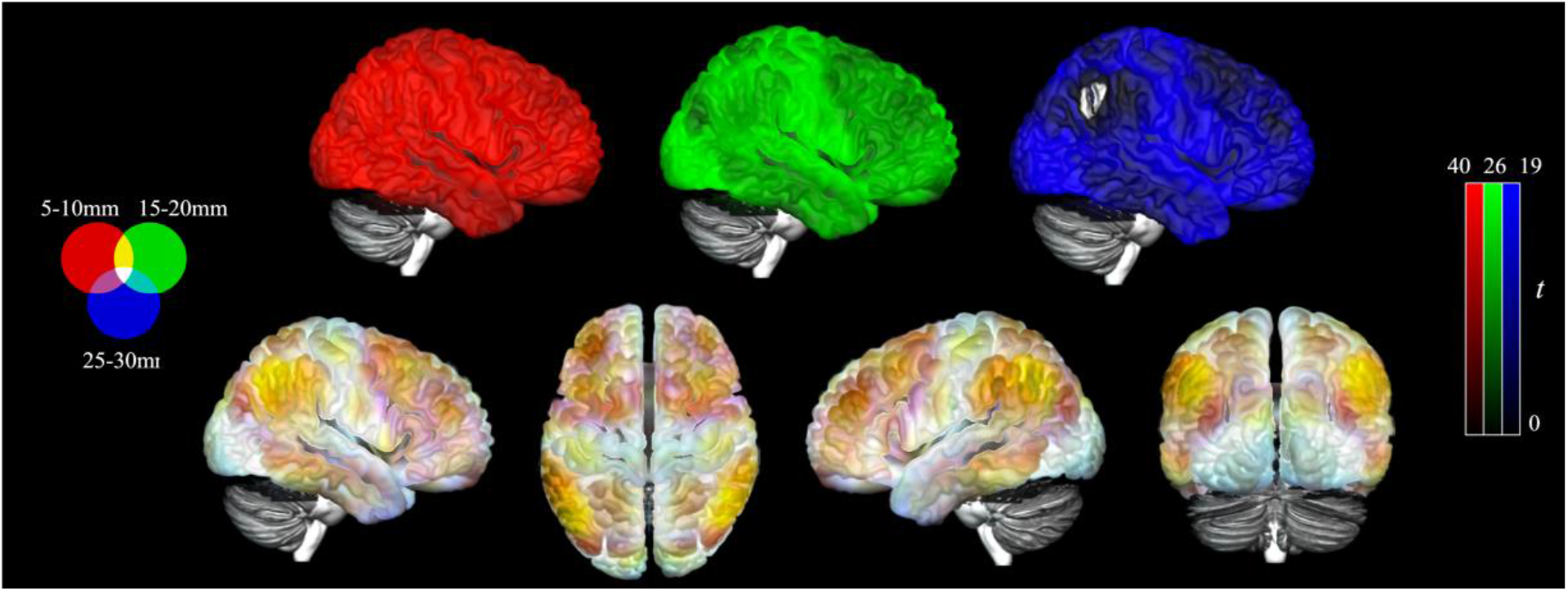
Iso-Distant Average Correlation (IDAC) brain maps across all three distances projected onto a cortical surface. Results displayed are from healthy control participants (*N* = 95) with overlay of the IDAC distances (top), 5-10 mm (red), 15-20mm (green), and 25-30mm (blue) with values capped at 95% of their respective maximal *t* values. The overlay of the 3 maps together (bottom) illustrates the consistent and unique effect across these distances through primary RGB colors and their secondary combinations. Left = Left.

### Group Differences in the Functional Structure of the Cerebral Cortex

The Effect of Group contrast demonstrated that MDD participants and healthy controls had significant differences across the three connectivity distances (Figure 2, Supplementary Table S1). These effects were most predominately observed in the bilateral posterior hippocampus extending to the retrosplenial cortex and ventral posterior cingulate as well as to the lingual and fusiform gyri. Additional large effects were observed in the dorsal mid to posterior insular cortices and right dorsal premotor cortex. Subsequent t-test for each distance 5–10, 15–20, and 25–30 mm illustrated that these results were all driven by greater local connectivity in MDD participants compared to controls (Figure 3). As such, across all distances MDD participants demonstrated greater local functional connectivity of the bilateral extended posterior hippocampus, fusiform and lingual gyri, dorsal mid-insula, and premotor cortex (Figure 4; Supplementary Table S2, S3 and S4).

**Figure 2.**
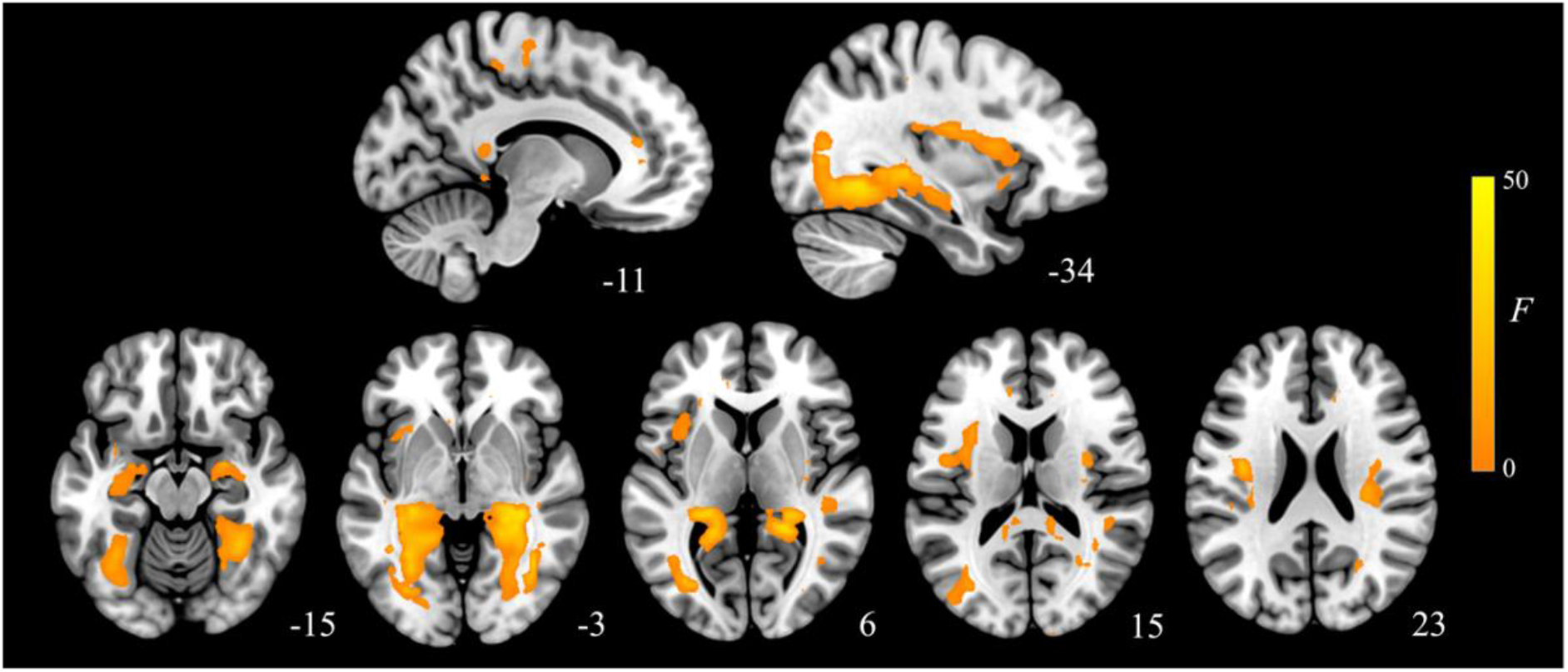
Main effects of group across the three local connectivity distances (5-10mm, 15-20mm, and 25-30mm). Results are displayed at *pFDR* < .008, whole-brain corrected.

**Figure 3.**
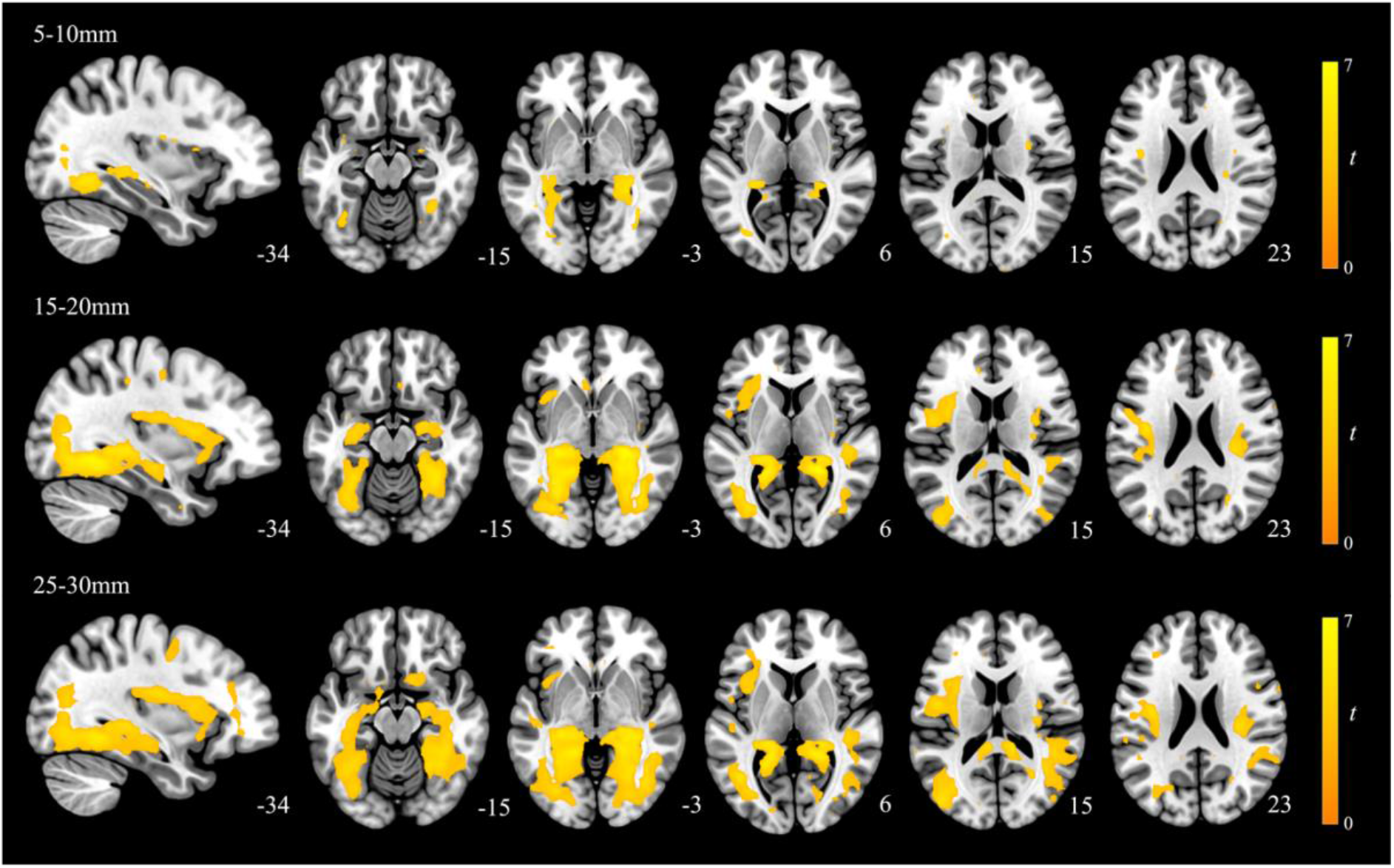
Group differences in the local functional connectivity for each of the 5-10mm, 15-20mm and 25-30mm distances (MDD participants > healthy controls). Results are displayed at *pFDR* < .008, whole-brain corrected.

**Figure 4.**
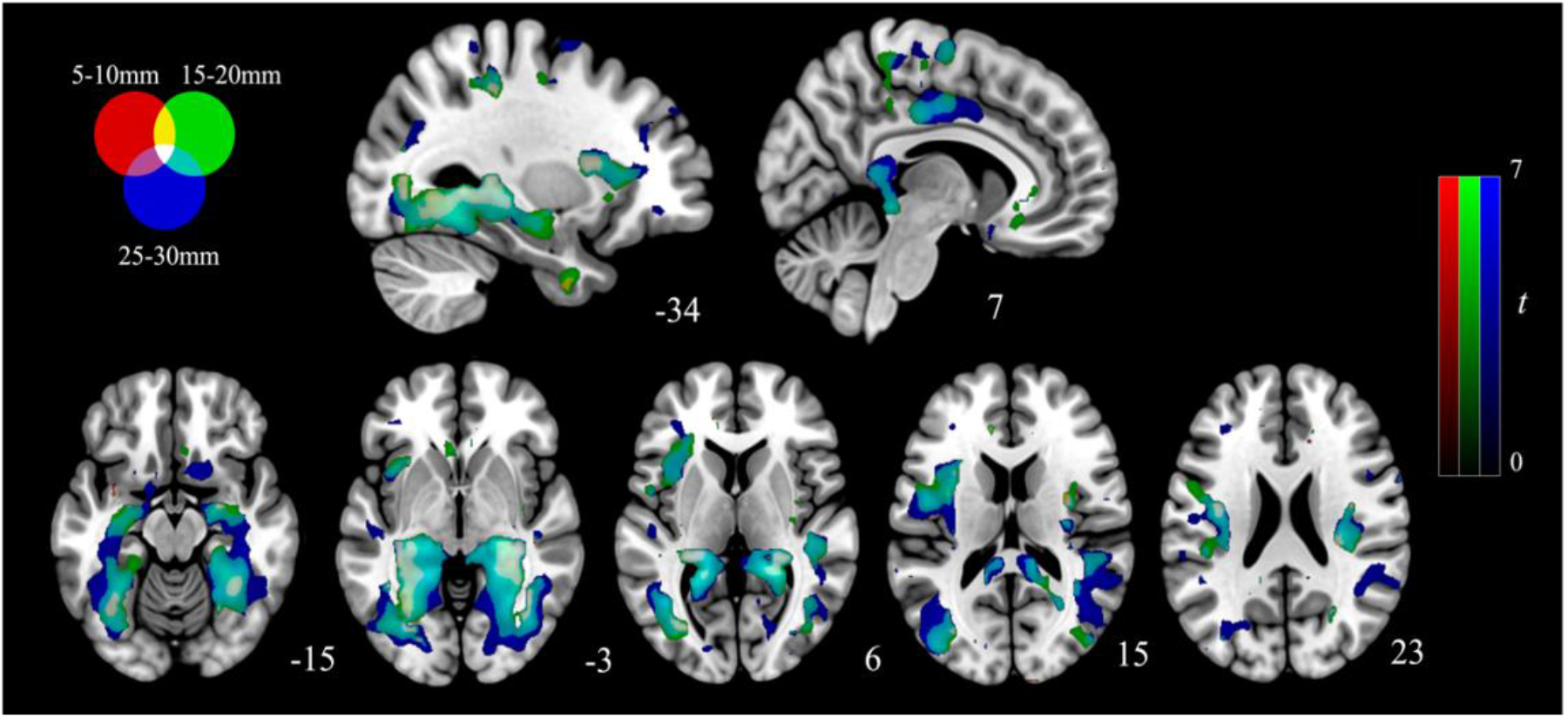
Red, green, and blue overlays illustrating the overlapping alterations to local functional connectivity present in MDD participants. Results are displayed at *pFDR* < .008, whole-brain corrected.

There were no significant group by distance interactions. However, the longer distances (15– 20 and 25–30 mm) demonstrated more apparent alterations in the left anterior insula, as well as across the cingulate, including the midcingulate (15–20 and 25–30 mm), ventral anterior (15–20 mm) and subgenual anterior cingulate (25–30 mm; Figure 3). The 25–30 mm distance was also observed to demonstrate differences between groups in the supplementary motor and premotor cortices.

### The Associations Between Depressive Symptoms, Emotional Regulation, and Local Functional Connectivity

We observed a significant association between ERQ Suppression scores and local connectivity across all three IDAC distances while adjusting for MADRS scores (Supplementary Figure S2). Specifically, there was a consistent association between these scores and the local connectivity of the bilateral mid-insula cortex for the short and medium lengths (5-10 mm and 15-20 mm). MADRS scores adjusted for ERQ suppression were conversely associated with connectivity alterations in the bilateral posterior hippocampi, lingual gyri, midcingulate and supplementary motor cortex across all three distanced (Supplementary Figure S3). Additional effects were observed in dorsal anterior cingulate (5-10 mm). No significant correlations were observed between IDAC measures and ERQ Reappraisal scores.

## Discussion

Our investigation of the differences in local functional connectivity patterns present in adolescents and young adults with MDD showed that MDD participants had greater multi-distance local functional connectivity across several brain regions, with the largest effects observed in the retrosplenial cortex, posterior hippocampus, dorsal mid-insula, and the fusiform and lingual gyri. We did not identify reduced local connectivity in the orbitofrontal cortex as hypothesized. Additionally, we observed consistent associations between ERQ Suppression scores and the local connectivity of regions including the insula across participants, which may have contributed to the between group differences observed in these areas.

### Posterior Hippocampal and Retrosplenial Alterations

The posterior hippocampus and retrosplenial cortex have consistently shown functional alterations in depression. Meta-analyses have identified increased regional homogeneity of the left hippocampus in drug naïve MDD participants (Hao *et al*., 2019). Similarly, reduced suppression of the hippocampus has been observed in depressed participants during loss reinforcement learning conditions, with the degree of this hyperactivity being associated with severity of symptoms (Johnston *et al*., 2015). Depression has also been associated with functional alterations in the retrosplenial cortex (Harel *et al*., 2016, Kumar *et al*., 2008, Zhu *et al*., 2012). Notably, both regions are components of the medial temporal subsystem of the DMN (Andrews-Hanna *et al*., 2010), and have been broadly implicated in affective memory and information processing, particularly autobiographical memory (Andrews-Hanna *et al*., 2014, Miller *et al*., 2014). The retrosplenial cortex has also been implicated in mental imagery and projecting oneself in time, both key components in episodic memory (Chrastil, 2018). As such, these regions are spatially and functionally positioned to link DMN subsystems necessary for episodic memory retrieval and core DMN regions, such as the posterior cingulate cortex, necessary for the generation of self-conceptualization (Andrews-Hanna *et al*., 2014, Davey and Harrison, 2022, Kaboodvand *et al*., 2018). Increased local synchrony of these regions observed in depressed individuals may therefore represent changes to the processing and encoding of emotional memory (Jaworska *et al*., 2015), and specifically contribute to the negatively biased recall of stimuli hypothesized by cognitive models of depression (Disner *et al*., 2011).

In rodent work the retrosplenial cortex has been implicated in abnormal long-term metabolic stress responses and vulnerability to depression (Harro *et al*., 2014). Excitatory efferent projections from the dorsal hippocampus, equivalent to the posterior hippocampus in humans (Fanselow and Dong, 2010), to the retrosplenial cortex have been identified as contributors to the generalization of negative memories (Ren *et al*., 2022). The hippocampus demonstrates a high proportion of glutamatergic pyramidal cells (Freund and Buzsaki, 1996, Olbrich and Braak, 1985), with co-activation of these neurons during rest being shown to significantly increase following chronic stress exposure (Tomar *et al*., 2021). Thus, due to the sensitivity of these regions to stress (Corcoran *et al*., 2018), hippocampus and retrosplenial cortex dysfunction may represent a potential mechanism through which early life adversity results in changes to the processing and encoding of emotional memory. This in turn, would facilitate the reductions in hippocampal volume observed in older depressed participants (Huang *et al*., 2013, MacQueen and Frodl, 2011, Malykhin and Coupland, 2015). It therefore appears that IDAC could be sensitive to stress-related neuroplastic changes impacting extended hippocampal function. Future examination of how these changes in local synchrony progress as a function of the chronicity of the disorder will provide insight about the longitudinal effect these local alterations have on other regions and networks.

### Dorsal Mid-Insula Alterations

Compared to healthy controls, MDD participants have shown decreased activity of the mid-insula cortex during interoceptive processing (Avery *et al*., 2014, DeVille *et al*., 2018) and recall (DeVille *et al*., 2018). MDD participants additionally illustrate increased resting-state connectivity between the dorsal mid-insula and affective regions, including the amygdala, subgenual cingulate cortex, and orbitofrontal cortex (Avery *et al*., 2014). It has been proposed that the insula illustrates a posterior-to-anterior progression of interoceptive processing, with the posterior insula mapping interoceptive signals (Craig, 2002, Kuehn *et al*., 2016), and the anterior insula associated with interoceptive awareness (Craig, 2009, Gu *et al*., 2013). Within this framework the dorsal mid-insula is well placed to serve as an intermediary between these processes. Primate cytoarchitectonics of the precentral insular gyrus further supports this functioning, as the mid-insula both projects to (Mesulam and Mufson, 1982), and receives input from (Mufson and Mesulam, 1982), the anterior and posterior insula. Moreover, the mid-insula has previously been associated with interoceptive processing (Centanni *et al*., 2021), including both interoceptive attention (Kurth *et al*., 2010, Schulz, 2016) and interoceptive recall (DeVille *et al*., 2018). Together this work indicates that this part of the insula may be responsible for processing interoceptive prediction errors, which occur following a mismatch between expectations concerning physiological states and bodily signals (Barrett and Simmons, 2015). A recent meta-analysis examining alterations across multiple psychiatric disorders during interoception identified consistent changes to the mid-insula (Nord *et al*., 2021). As such, the integrative role of the mid-insula may increase its sensitivity to dysfunctional signals emerging from disparate bodily and brain systems (Nord *et al*., 2021, Sliz and Hayley, 2012) thereby contributing to abnormal emotional states (Namkung *et al*., 2018).

Interestingly, we observed that the local functional connectivity of the insula was associated with habitual use of expressive suppression. Due to expressive suppression involving inhibiting the outward expression of emotional behavior (Gross, 2015), its use may be viewed as an attempt to exert top-down control of interoceptive signals (Nord *et al*., 2021, Paulus and Stein, 2010). Maladaptive emotional regulation strategies are well noted in depressive samples (Schafer *et al*., 2017), including increased use of expressive suppression and decreased use of cognitive reappraisal (Joormann and Gotlib, 2010). While a recent review has highlighted mixed evidence concerning whether expressive suppression is associated with depression (Dryman and Heimberg, 2018), longitudinal research suggests that expressive suppression may both predict (Larsen *et al*., 2013) and be predicted by depressive symptoms (Larsen *et al*., 2012). The precise nature of this relationship likely depends on the developmental stage being investigated (Dryman and Heimberg, 2018). Age has been suggested as an important mediator of this relationship given that older adults use expressive suppression more frequently, however, experience less psychological distress from its use (Brummer *et al*., 2014). Thus, the young nature of our sample may explain the large difference in expressive suppression observed between the healthy controls and MDD participants.

### Limitations

The strength and novelty of this study should be considered in the context of its limitations. Our results were derived from young adults and adolescents. It remains unclear whether these same efects would be present in older populations and how these local connectivity profiles change longitudinally and as a function of illness chronicity. Additionally, while the homogeneity of the current sample is a major strength it also limits the generalizability of our findings. As such, further work in transdiagnostic samples examining the relationship between local functional connectivity and factors such as expressive suppression may be warranted.

## Conclusion

We have examined the local functional connectivity of the cerebral cortex using a novel approach to investigate brain alterations associated with MDD. We identified differences in the local synchrony of several regions, including the retrosplenial cortex, posterior hippocampus, and dorsal mid-insula. Across these regions depressed individuals demonstrated increased coupling of intracortical activity. Additionally, across participants, increased local functional connectivity of the insula was correlated with use of expressive suppression as an emotional regulation strategy. Longitudinal examination of these between group local synchrony differences will aid in identifying whether this relationship changes as a function of the chronicity of the disorder.

## Acknowledgments

The authors thank Katerina Stephanou, Lisa Incerti and Rebecca Kerestes for contributions to data collection, as well as staff from the Sunshine Hospital Medical Imaging Department (Western Health, Melbourne).

