## Supplementary Materials for "Graded changes in local functional connectivity of the cerebral cortex in young people with depression"

**Supplementary Tables………………………………………………………………………2**

Supplementary Table S1: Group differences in Iso-Distance Average Correlation (IDAC) functional connectivity measures across the three distances (F contrast Effect of Group).….2

Supplementary Table S2: Group differences in functional connectivity measures based on Iso-Distance Average Correlations (IDAC) for the 5-10mm distance………..……………...4

Supplementary Table S3: Group differences in functional connectivity measures based on Iso-Distance Average Correlations (IDAC) for the 15-20mm distance………..…………….5

Supplementary Table S4: Group differences in functional connectivity measures based on Iso-Distance Average Correlations (IDAC) for the 25-30mm distance………..…………….7

**Supplementary Figures……………………………………………………………………..9**

Supplementary Figure S1: Distribution of (A) Montgomery-Åsberg Depression Rating Scale scores, (B) Emotion Regulation Questionnaire Suppression Subscale scores and (C) Emotion Regulation Questionnaire Reappraisal Subscale scores by diagnostic group. Healthy controls are shown in blue, MDD participants in green.………..…………………..………………...9

Supplementary Figure S2: Red, green, and blue overlays illustrating associations between ERQ Suppression scores and local functional connectivity.……………………………….10

Supplementary Figure S3: Red, green, and blue overlays illustrating associations between MADRS scores and local functional connectivity.……………………………………..….11

| **Supplementary Tables**  Supplementary Table S1  *Group differences in Iso-Distance Average Correlation (IDAC) functional connectivity measures across the three distances (F contrast Effect of Group)* | | | | | | | |
| --- | --- | --- | --- | --- | --- | --- | --- |
| Brain region | Hemisphere | BA | Coordinates | | | Cluster size (3mm^3^ voxels) | *F*-value |
|  |  |  | X | Y | Z |  |  |
| *MDD > Controls* |  |  |  |  |  |  |  |
| Hippocampus | R | - | 31 | -39 | 0 | 882 | 46.57 |
| Retrosplenial cortex | R | 30 | 22 | -42 | 3 |  | 41.60 |
| Hippocampus | R | - | 28 | -30 | -3 |  | 41.29 |
| Fusiform gyrus | L | 37 | -35 | -57 | -6 | 883 | 45.54 |
| Hippocampus | L | - | -26 | -36 | 9 |  | 38.50 |
|  |  | - | -32 | -39 | 0 |  | 37.95 |
| Insula | L | 13 | 34 | -3 | 18 | 78 | 32.62 |
|  |  | 13 | 31 | -15 | 18 |  | 27.63 |
| Supramarginal gyrus | L | 40 | 37 | -30 | 24 |  | 26.55 |
| Primary motor cortex | L | 4 | -38 | -9 | 21 | 237 | 32.21 |
| Insula | L | 13 | -35 | 0 | 18 |  | 28.56 |
| Supramarginal gyrus | L | 40 | -44 | -30 | 27 |  | 24.80 |
| Supplementary motor cortex | L | 6 | -14 | -15 | 63 | 15 | 28.44 |
| Primary motor cortex | L | 4 | -11 | -21 | 57 |  | 24.12 |
| Sensory association cortex | R | 5 | -17 | -30 | 48 | 32 | 27.40 |
|  | L | 5 | -17 | -36 | 42 |  | 26.13 |
| Precuneus | L | 7 | -14 | -45 | 60 |  | 15.80 |
| Angular gyrus | R | 39 | 37 | -51 | 18 | 10 | 26.83 |
|  | R |  | 40 | -57 | 12 |  | 15.85 |
| Supplementary motor cortex | R | 6 | 16 | -21 | 42 | 121 | 25.54 |
| Ventral anterior cingulate | R | 24 | 13 | -9 | 42 |  | 23.50 |
| Supplementary motor cortex | R | 6 | 1 | -12 | 69 |  | 23.15 |

| Supplementary Table S2  *Group differences in functional connectivity measures based on Iso-Distance Average Correlations (IDAC) for the 5-10mm distance* | | | | | | | |
| --- | --- | --- | --- | --- | --- | --- | --- |
| Brain region | Hemisphere | BA | Coordinates | | | Cluster size (3mm^3^ voxels) | t-value |
|  |  |  | X | Y | Z |  |  |
| *MDD > Controls* |  |  |  |  |  |  |  |
| Hippocampus | R | - | 31 | -39 | 0 | 219 | 5.96 |
| Visual association cortex | R | 19 | 40 | -60 | -3 |  | 5.53 |
| Caudate tail | R | - | 34 | -30 | -3 |  | 5.50 |
| Fusiform gyrus | L | 37 | -35 | -57 | -6 | 205 | 5.89 |
| Caudate tail | L | - | -26 | -36 | 9 |  | 5.34 |
|  | L | - | -35 | -33 | 0 |  | 5.28 |
| Insula | R | 13 | 34 | -3 | 18 | 24 | 5.43 |
| Primary motor cortex | L | 4 | -38 | -6 | 24 | 18 | 5.91 |
| Supplementary motor area | L | 6 | -35 | 0 | 18 |  | 4.36 |
| Insula | L | 13 | -29 | 9 | 12 | 13 | 4.85 |
| Pars opercularis | L | 44 | -38 | 12 | 12 |  | 3.89 |

| Supplementary Table S3  *Group differences in functional connectivity measures based on Iso-Distance Average Correlations (IDAC) for the 15-20mm distance* | | | | | | | |
| --- | --- | --- | --- | --- | --- | --- | --- |
| Brain region | Hemisphere | BA | Coordinates | | | Cluster size (3mm^3^ voxels) | t-value |
|  |  |  | X | Y | Z |  |  |
| *MDD > Controls* |  |  |  |  |  |  |  |
| Retrosplenial cortex | R | 30 | 22 | -45 | 6 | 1228 | 6.96 |
| Hippocampus | R | - | 28 | -30 | -3 |  | 6.91 |
|  | R | - | 28 | -39 | 0 |  | 6.58 |
| Visual association cortex | L | 19 | -32 | -54 | -6 | 1239 | 6.74 |
| Hippocampus | L | - | -29 | -30 | -3 |  | 6.52 |
| Retrosplenial cortex | L | 30 | -17 | -45 | 3 |  | 6.52 |
| Primary motor cortex | L | 6 | -38 | -9 | 24 | 406 | 5.57 |
| Insula | L | 13 | -35 | 0 | 18 |  | 5.38 |
| Premotor cortex | L | 6 | -44 | -3 | 18 |  | 5.28 |
| Insula | R | 13 | 31 | -15 | 18 | 196 | 5.34 |
|  | R | 13 | 31 | -6 | 12 |  | 5.11 |
| Supramarginal gyrus | R | 40 | 37 | -30 | 24 |  | 5.04 |
| Mid cingulate cortex | R | 24 | 16 | -18 | 42 | 347 | 5.23 |
|  | R | 24 | 13 | -9 | 42 |  | 4.88 |
| Supplementary motor cortex | L | 40 | -14 | -12 | 48 |  | 4.75 |
| Sensory association cortex | L | 5 | -17 | -30 | 48 | 67 | 4.88 |
|  | L | 5 | -14 | -36 | 54 |  | 4.80 |
|  | L | 5 | -17 | -36 | 42 |  | 4.33 |
| Primary motor | L | 24 | -11 | -21 | 57 | 16 | 4.71 |
| Supplementary motor cortex | L | 5 | -14 | -15 | 63 |  | 4.71 |
| Premotor cortex | L | 6 | -29 | -12 | 48 | 12 | 4.38 |
| Mid cingulate cortex | L | 24 | -8 | 3 | 36 | 22 | 4.35 |
| Subgenual anterior cingulate cortex | L | 25 | -5 | 24 | -3 | 15 | 4.33 |
| Rostral anterior cingulate | R | 24 | 1 | 18 | 0 |  | 3.84 |
| Ventral posterior cingulate | L | 23 | -8 | -18 | 33 | 11 | 4.26 |
| Sensory association cortex | L | 5 | -26 | -36 | 45 | 14 | 4.03 |
| Dorsal anterior cingulate | R | 32 | 13 | 3 | 39 | 11 | 3.95 |
|  | R | 32 | 13 | 12 | 39 |  | 3.73 |
| Subgenual anterior cingulate cortex | R | 25 | 4 | 24 | -12 | 10 | 3.93 |
| Supplementary motor cortex | L | 6 | -20 | -6 | 66 | 12 | 3.53 |
|  | L | 6 | -11 | -6 | 69 |  | 3.53 |

| Supplementary Table S4  *Group differences in functional connectivity measures based on Iso-Distance Average Correlations (IDAC) for the 25-30mm distance* | | | | | | | |
| --- | --- | --- | --- | --- | --- | --- | --- |
| Brain region | Hemisphere | BA | Coordinates | | | Cluster size (3mm^3^ voxels) | t-value |
|  |  |  | X | Y | Z |  |  |
| *MDD > Controls* |  |  |  |  |  |  |  |
| Retrosplenial cortex | L | 30 | -5 | -39 | 12 | 1530 | 6.68 |
|  | L | 30 | -17 | -45 | 0 |  | 6.46 |
| Fusiform gyrus | L | 37 | -38 | -51 | -9 |  | 6.31 |
| Retrosplenial cortex | R | 30 | 22 | -42 | 3 | 1937 | 6.46 |
| Visual association cortex | R | 19 | 25 | -54 | -9 |  | 5.97 |
| Parahippocampal cortex | R | 36 | 37 | -39 | -9 |  | 5.95 |
| Primary motor cortex | L | 4 | -35 | -12 | 21 | 485 | 5.53 |
| Premotor cortex | L | 5 | -47 | -3 | 18 |  | 5.26 |
| Insula | L | 13 | -35 | 0 | 18 |  | 4.81 |
| Ventral posterior cingulate cortex | R | 23 | 10 | -18 | 36 | 808 | 5.20 |
| Mid cingulate cortex | R | 24 | 16 | -21 | 42 |  | 5.12 |
| Premotor cortex | R | 6 | 34 | -6 | 45 |  | 4.82 |
| Primary sensory cortex | R | 1 | 52 | -21 | 57 | 63 | 4.67 |
|  | R | 1 | 34 | -27 | 42 |  | 3.59 |
|  | R | 1 | 40 | -21 | 36 |  | 3.55 |
| Subgenual anterior cingulate cortex | R | 25 | 10 | 12 | -18 | 54 | 4.54 |
| Putamen | R | - | 25 | 12 | -12 |  | 4.19 |
| Insula | R | 13 | 37 | 12 | -12 |  | 3.61 |
| Superior temporal gyrus | L | 22 | -47 | -21 | -6 | 29 | 4.53 |
| Angular gyrus | L | 39 | -44 | -45 | 30 | 13 | 4.04 |
| Temporal pole | R | 38 | 40 | 18 | -36 | 11 | 3.89 |
| Dorsal posterior cingulate cortex | R | 31 | 25 | -63 | 27 | 18 | 3.84 |
| Dorsal posterior cingulate cortex | L | 31 | -17 | -36 | 42 | 20 | 3.81 |
| Sensory association cortex | L | 5 | -17 | -30 | 48 |  | 3.79 |
|  | L | 5 | -11 | -27 | 54 |  | 3.51 |
| Primary visual cortex | L | 17 | -17 | -87 | 6 | 11 | 3.68 |
| Primary motor cortex | R | 4 | 10 | -24 | 66 | 10 | 3.60 |
|  | R | 4 | 4 | -33 | 63 | 14 | 3.60 |

Supplementary Figures

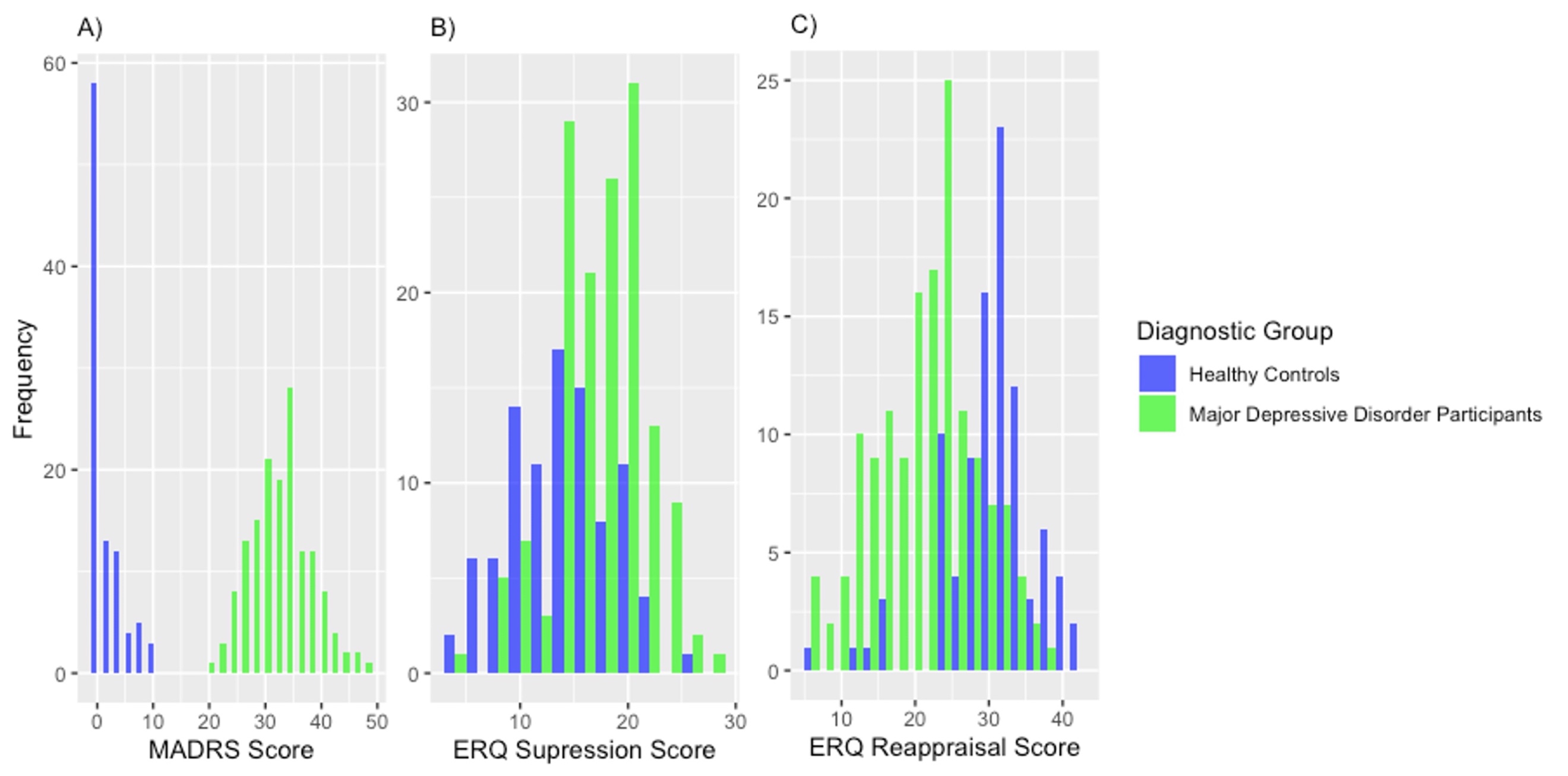

*Supplementary Figure S1.* Distribution of (A) Montgomery-Åsberg Depression Rating Scale scores, (B) Emotion Regulation Questionnaire Suppression Subscale scores and (C) Emotion Regulation Questionnaire Reappraisal Subscale scores by diagnostic group. Healthy controls are shown in blue, MDD participants in green.

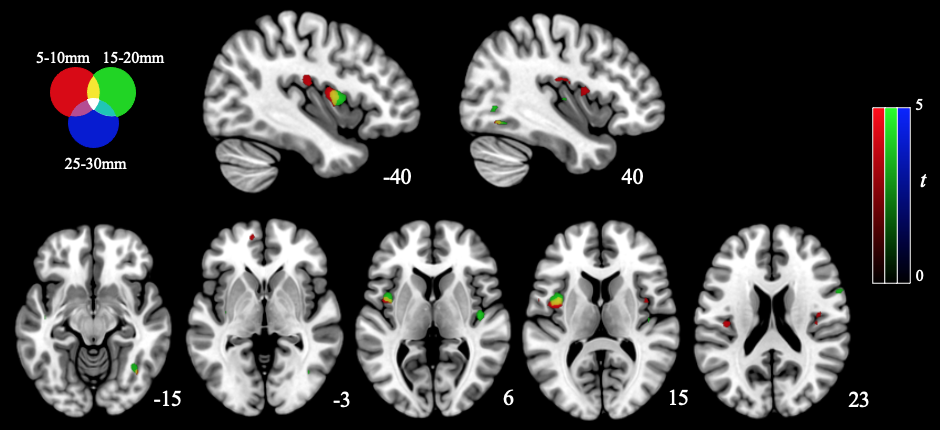

*Supplementary Figure S2.* Red, green, and blue overlays illustrating associations between ERQ Suppression scores and local functional connectivity. Results are displayed at p < .001, family-wise error cluster corrected.

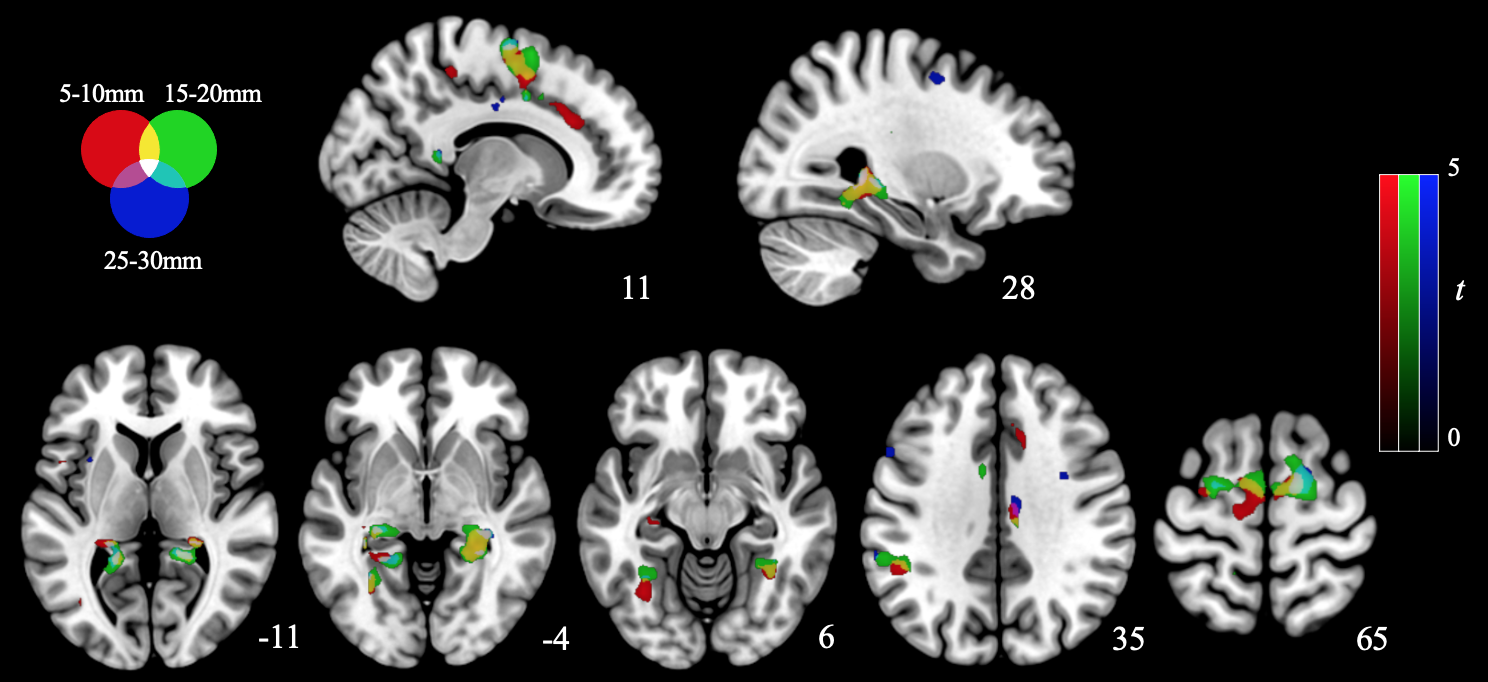

*Supplementary Figure S3.* Red, green, and blue overlays illustrating associations between MADRS scores and local functional connectivity. Results are displayed at *p* < .001, family-wise error cluster corrected.
